# *Ripply3* overdosage induces mid-face shortening through *Tbx1* downregulation in Down syndrome models

**DOI:** 10.1101/2024.09.13.612914

**Authors:** José Tomás Ahumada Saavedra, Claire Chevalier, Agnes Bloch Zupan, Yann Herault

## Abstract

The most frequent and unique features of Down syndrome (DS) are learning disability and ucraniofacial (CF) dysmorphism. The DS-specific CF features are an overall reduction in head dimensions (microcephaly), relatively wide neurocranium (brachycephaly), reduced mediolaterally orbital region, reduced bizygomatic breadth, small maxilla, small mandible, and increased individual variability. Until now, the cellular and molecular mechanisms underlying the specific craniofacial phenotype have remained poorly understood. Investigating a new panel of DS mouse models with different segmental duplications on mouse chromosome 16 in the region homologous to human chromosome 21, we identified new regions and the role of two candidate gene for DS-specific CF phenotypes. First, we confirmed the role of *Dyrk1a* in the neurocranium brachycephaly. Then, we identified the role of the transcription factor *Ripply3* overdosage in midface shortening through the downregulation of *Tbx1*, another transcription factor involved in the CF midface phenotype encountered in DiGeorge syndrome. This last effect occurs during branchial arches development through a reduction in cell proliferation. Our findings define a new dosage-sensitive gene responsible for the DS craniofacial features and propose new models for rescuing all aspects of DS CF phenotypes. This data may also provide insights into specific brain and cardiovascular phenotypes observed in DiGeorge and DS models, opening avenues for potential targeted treatment to soften craniofacial dysmorphism in Down syndrome.

## INTRODUCTION

Trisomy 21, or Down syndrome (DS), is a pleiotropic disorder with intellectual disability, CF changes, and comorbidities. Somehow, gene dosage effects of one or more of the 671 genes on human chromosome 21 (Hsa21) are responsible for specific pathological features (Zhu et al. 2019; Duchon and Herault 2016). Facial features are characteristic of individuals with DS (Kisling 1966), although their severity varies from one individual to another (Roper and Reeves 2006). DS-related CF dysmorphism includes an overall reduction in head dimensions (microcephaly), relatively wide neurocranium (brachycephaly), small midface, reduced mediolaterally orbital region, reduced bizygomatic breadth, small maxilla, and small mandible (Fink, Madaus, and Walker 1975; Farkas, Kolar, and Munro 1985; Allanson et al. 1993; Baxter et al. 2000). Patients also experience a low bone mass associated with reduced osteoblast activity and high bone turnover (McKelvey et al. 2013).

Studies in rodents and humans have attempted to identify the candidate gene(s) causing DS clinical features (Korbel et al. 2009; McCormick et al. 1989). Using the rapid engineering of the *Mus musculus* (Mmu) genome, multiple DS mouse models have been generated (Dierssen, Herault, and Estivill 2009; Herault et al. 2017) containing extra copies of the Hsa21-orthologous regions of three murine chromosomes: Mmu chromosome 16 (Mmu16), 10 (Mmu10), and 17 (Mmu17) (Reeves et al. 1995; Sago et al. 1998).

DS mouse models have previously been studied for CF phenotypes. The most notable studies have examined the Ts(17^16^)65Dn model, hereafter termed Ts65Dn. This mouse strain carries an extra mini-chromosome with the *mIR155-Zbtb21* region of Mmu16 translocated downstream of *Pde10a,* close to the centromere of Mmu17 (Duchon et al. 2011). Thus, Ts65Dn is trisomic for 104 of the Hsa21 orthologs of the Mmu16 between *miR155 and Zbtb21* (Olson et al. 2004; Gardiner et al. 2003; Muñiz Moreno et al. 2020). The Ts65Dn model displays a variety of phenotypes similar to those found in DS individuals (Olson et al. 2004), including a low bone mass caused by intrinsic cellular defects in osteoblast differentiation, reducing bone formation (Fowler et al. 2012). In addition, bone resorption mediated by osteoclasts is also reduced, but this is not enough to overcome the low rate of bone formation (Parsons et al. 2007; Thomas et al. 2021). These animals also show many cognitive and behavioral traits as well as characteristic skeletal, craniofacial, cardiovascular features, granule-cell density of the dentate gyrus, and megakaryocytopoiesis mimicking the phenotype encountered in individuals with Down syndrome (Baxter et al. 2000). The CF phenotypes found include brachycephaly, reduced facial and cranial vault dimension, reduced cerebellar volume, and many features present in individuals with DS. CF changes even more similar to those in humans were found in our new Ts(17^16^)66Yah model, devoid of non-Hsa21 triplicated genes (Duchon et al. 2022), confirming a significant contribution of one or more genes found between *mIR155* and *Zbtb21* to the CF phenotypes.

CF defects have also been detected in other DS mouse models. The Ts(16C-tel)1Cje (Ts1Cje) model carries a translocation that encompasses 81 orthologous genes between *Sod1* and *Mx1* (Sago et al. 1998; Huang et al. 1997) and displays a generalized reduction in CF size with additional features (Richtsmeier et al. 2002). By contrast, the Dp(16*Cbr1-Fam3b*)1Rhr (noted here Dp1Rhr), a model trisomic for 33 genes (Olson et al. 2007), exhibited a larger overall size and CF alterations, including more pronounced defects in the mandible than observed in Ts65Dn mice and individuals with DS (Deitz and Roper 2011).

The Dp(16*Lipi-Zbtb21*)1Yey mouse model (Dp1Yey) is a larger model with a 22.9 Mb direct duplication of the entire Mmu16 region in conserved synteny with Hsa21, containing 118 orthologous protein-coding genes (Li et al. 2007). The CF phenotype corresponds to brachycephaly, a reduced dimension of the maxillary and palate, and reduced mandibular size. The skulls also exhibited increased variance relative to euploid littermates for specific linear distances (Li et al. 2007; Yu et al. 2010; Starbuck et al. 2014).

Another model quite similar to Dp(16)1Yey, but with a slightly different duplicated interval, is the Dp(16*Lipi-Zbtb21*)1TybEmcf, or Dp(16)1Tyb (Lana-Elola et al. 2016). In 2023, using morphometric analysis of the Dp1Tyb mouse model of DS and an associated mouse genetic mapping panel, Redhead et al. (Redhead et al. 2023) showed that *Dyrk1a* is required in three copies to cause CF dysmorphology in Dp(16)1Tyb mice. In addition, Dp(16)1Tyb mice display many phenotypic features characteristic of DS in humans, including congenital heart defects, reduced bone density, and deficits in memory, locomotion, hearing, and sleep (Chang et al. 2020; Lana-Elola et al. 2011; 2016; Thomas et al. 2020; Watson-Scales 2018). Taken together, the candidate genes responsible for craniofacial phenotypes found in DS models include *Dyrk1a*, *Rcan1* (*Dscr1*), and *Ets2*. *Dyrk1a* has been implicated in several DS phenotypes, including cognitive impairment, motor function, and craniofacial abnormalities (Hämmerle et al. 2003; Arron et al. 2006; Atas-Ozcan et al. 2021). Jhonson et al. in 2024, showed that a decreased in *Dyrk1a* in Xenopus resulted in craniofacial malformations, altered expression of critical craniofacial regulators as *Pax3* and *Sox9* fundamental for cranial neural crest development, and presented altered retinoic acid, hedgehog, nuclear factor of activated T cells (*NFAT*), *Notch* and *WNT* signaling pathways. These results indicate that DYRK1A function is critical for early craniofacial development and must properly regulate the expression of specific craniofacial regulators in the branchial arches (Johnson et al. 2024).

Disruption of *Tbx1* expression is a common aspect of CF dysmorphias. *Tbx1* Is the first dosage-sensitive gene identified in the DiGeorge syndrome (DGS)/velocardiofacial syndrome (VCFS), a congenital disorder characterized by neural-crest-related developmental defects. In humans, *TBX1* haploinsufficiency causes craniofacial anomalies (Lindsay et al. 2001). In the mouse model for DiGeorge syndrome the phenotype observed in the mutant mice for the T-box gene, *Tbx1*^+/-^, encompasses abnormal development of the skeletal structures derived from the first and second pharyngeal arches, with reduced dimension of the midface (Jerome and Papaioannou 2001); a similar situation found in the DS mouse models.

However, the details of how the dosage imbalance of Hsa21 genes affects CF morphogenesis are still poorly understood (Starbuck et al. 2017). Unraveling the complex genetics and adaptative biological processes involved in forming craniofacial structures is essential. Many genes are conserved across mammals, implying that the genetic programs for a specific phenotype may also be conserved. Therefore, this can validate the study of animal models to decipher human genetic outcomes (Richtsmeier, Baxter, and Reeves 2000).

As observed in different models, human partial trisomy has allowed mapping areas of Hsa21 to contribute to craniofacial anomalies, but a specific region has not yet been identified (Lyle et al. 2009; Korbel et al. 2009). Identifying the dosage-sensitive genes responsible for each element of the DS phenotype will help us better understand the molecular mechanisms underlying the various symptoms and will allow us to define therapeutic options better (Antonarakis 2017; Antonarakis et al. 2020).

The Cre-LoxP technology has enabled the engineering of more precise duplications (Hérault et al. 2010; Ruf et al. 2011). Applying this technology, we generated mouse models carrying different segmental duplications of regions located on the Mmu16 homologous to Hsa21. In this study, we used these new DS models and two already known models, Dp(16)1Yey and Tg(Dyrk1a), to establish correlations between human-related CF phenotype and genotype and to understand the potential craniofacial effect of the duplication of different chromosomal regions via a morphometric analysis of the animal models. This led us to narrow our research to find new Mmu16 regions involved in CF and identify corresponding candidate genes responsible for the DS-CF phenotype.

## MATERIALS AND METHODS

### Previously reported rodent models used

The Dp(16)1Yey and Tg(*Dyrk1a*) (official name Tg(*Dyrk1a*)189N3Yah) models (Li et al. 2007; Guedj et al. 2012) were maintained on the C57BL/6J genetic background. We also used the SD:CRL Dp (11Lipi-Zbtb21)1Yah (short name Dp(Rno11)) rat model generated in the lab (Birling et al. 2017) that carries a duplication of the *Lipi-Zbtb21*, an interval similar to the mouse Dp(16)1Yey, found on rat chromosome 11.

### Generation of the new DS mouse strains

These new lines were generated via an *in vivo* chromosomal recombination technique, which combines a transposon system (Ruf et al. 2011) and a meiotic recombination system Cre-LoxP (Hérault et al. 1998). The transposon system consists of the transposase enzyme and its substrate, the transposon. The enzyme recognizes specific repeat sequences (ITR) flanked on either side of a given DNA sequence (in this case, a vector containing a specific *loxP* site) (Ruf et al. 2011). Once a region of interest is bounded by two *loxP* sites, a transgene expressing the Cre recombinase is brought into the same individual by successive crosses. In this animal, the Cre enzyme recombines the sequences of the *loxP* sites to produce a duplication (or partial trisomy) of the region of interest (Hérault et al. 1998; Hérault et al. 2010).

The new mouse models have been developed with the following segmental duplications in the Mmu16. For Dp(16*Samsn1-Cldn17*)7Yah (Dp(16)7Yah) we duplicated the segment between *Samsn1* and *Cldn17*. Dp(16*Tiam1-Clic6*))8Yah (Dp(16)8Yah) presents a duplication between *Tiam1* and *Clic6*. Dp(16*Cldn17-Brwd1*))9Yah (Dp(16)9Yah) displays a duplication in the interval between *Cldn17* and *Brwd1*. Dp(16*Tmprss15-Setd4*)10Yah (Dp(16)10Yah) has the segment between *Tmprss15* and *Setd4* duplicated, similar to Dp(16*Tmprss15-Grik1*)11Yah (Dp(16)11Yah), but this model presents a region duplicated until *Grik1*. Dp(16*Tmprss15-Zbtb21*)12Yah (Dp(16)12Yah) has the duplicated region from *Tmprss15* to *Zfp295*, and Dp(16*Cldn17-Vps26c*(*Dyrk1a*KO))13Yah (Dp(16)13Yah) from *Cldn17* to *Vps26c*, up to the sequence of *Dyrk1a* which is inactivated. All lines were maintained on C57BL/6J genetic background (Fig. 2A).

### Adult cohorts generated

Mice were housed under specific pathogen-free (SPF) conditions and were treated in compliance with the animal welfare policies of the French Ministry of Agriculture (law 87 848). As a major genotype effect compared to sex was previously described elsewhere independently (Redhead et al. 2023), we decided to use females. For each mouse line, about ten littermates by each genotype, DS, and wild-type (WT) were collected (n = 180). We tried to have balanced males and females in the cohorts. For example, For the Dp(16)1Yey line, six females plus five males for the dup carrier and six males plus three females for control were used. Nevertheless, this was not the case in all the other lines, with sometimes more female individuals collected than males, because males were used to breed the lines.

### Micro-computed tomography scan of the skull of mutant and control mouse lines

Animals were euthanized with the standard procedure at 14 weeks old. Briefly, the mouse heads were dissected apart from the body. A polystyrene section was interposed between the mandible and maxilla to separate the jaws. After dissection, samples were fixed in a 4% paraformaldehyde solution (PFA), washed with water, and stored in 70% ethanol. The mouse heads were scanned using the Quantum FX micro-computed tomography imaging system (Caliper Life Sciences, Hopkinton, MA, USA) to evaluate the morphology of the skull and mandible. The images obtained were delivered in DICOM format. The scan parameters used to carry out the scanning of the samples correspond to 2 scans of every sample, anterior part, and posterior part using the mode Scan Technique Fine of 2 minutes, with a field of view (FOV) of 40 mm, the voltage 90 kV, CT 160 μA, resolution pixel size 10 µm and the capture size for live mode viewing in small, live current 80kV.

### Imaging Processing

For each sample, two scans were obtained, one from the anterior area of the skull and one from the posterior region. FIJI software was used to unite these two scans and create a single file, performing the plugin “Stitching” and making one file in TIFF format. This format can be opened using different image processors. Stratovan Checkpoint software (Stratovan Corporation, Sacramento, USA, Version 2018.08.07. Aug 07, 2018.) was used to place the landmarks (Table S1 and S2, supplementary information), extract the 3D coordinates, create Procrustes average models, and perform the voxel analysis. 3dMD Vultus® software (3dMD LLC, Atlanta, GA, USA) generated heat maps.

### Morphometrics analysis

Morphometrics is the quantification and statistical analysis of form. Form is the combination of size and shape of a geometric object in an arbitrary orientation and location (shape is what remains of the geometry of such an object once it is standardized for size). Various approaches can be employed when conducting morphometric analysis. The method of interest in this study is the landmark-based method, which is a conventional approach that relies on phenotypic measurements such as linear distances, angles, weights, and areas. In this case, we used 61 landmarks, 39 in the skull and 22 in the mandible, to obtain the 3D coordinates of the structure (Hallgrimsson et al. 2015).

Based on 3D coordinates, Euclidean Distance Matrix Analysis (EDMA) is one of the principal tools for analyzing landmark-based morphometric data (Lele and Richtsmeier 2001). This method builds a matrix of linear distances between all possible pairs of landmarks for each specimen (Lele et Richtsmeier 1991). Morphological differences between groups can be pinpointed to specific linear distances on an object through pairwise comparisons of mean form or shape matrices, followed by bootstrapping to estimate the significance of these differences (Lele and Richtsmeier 2001). In this study, two tests were done for each group of samples, first to analyze the form of the skull and mandibles with form difference matrix (FDM) and then the shape with the shape difference matrix (SDM).

In addition, to track the landmarks associated with a significant change and understand where they are located in the CF structures, “EDMA FORM or SHAPE Influence landmark analysis” was performed (Cole and Richtsmeier 1998). The purpose of this test is to search which landmarks present a Relative Euclidean distance (RED) > 1.05 or < 0.95 (outside of the confidence interval 97,8%), meaning, which landmarks show a bigger difference in linear distances between every landmark and in what direction.

Another way to handle landmark-based data is using a multivariate statistical analysis of form, geometric morphometric. This method relies on the superimposition of landmark coordinate data to place individuals into a common morpho-space. The most used superimposition form is the Generalized Procrustes (GP) method and Principal Component Analysis (PCA). This method places multiple individual specimens into the same shape space by scaling, translating, and rotating the landmark coordinates around the centroid of every sample (Rohlf and Slice 1990). As an alternative, we took advantage of Stratovan Checkpoint (Stratovan Corporation, Sacramento, USA) to create population average models and perform a voxel-based analysis, where we can observe directly in 3D models the changes between populations.

Finally, using the 3dMD Vultus® software, we created Procrustes average models created in Checkpoint to perform a landmarking calculation.

### Whole-Mount Skeletal Staining

DS individuals also experience a low bone mass associated with reduced osteoblast activity and bone turnover (McKelvey et al. 2013). To study these defects in ossification, we performed a skeletal/cartilage staining with alizarin red and alcian blue in a representative DS mouse model, Dp(16)1Yey.

Whole-mount skeletal staining permits the evaluation of the shapes and sizes of skeletal elements. Thus, it is the principal method for detecting changes in skeletal patterning and ossification. Because cartilage and bone can be distinguished by differential staining, this technique is also a powerful means to assess the pace of skeletal maturation.

We collected n=20 specimens, ten samples (5 WT vs 5 Dp(16)1yey) for the embryonic stage (E) 18.5 and 10 samples (5 WT vs 5 Dp(16)1yey) for P2 (2 days after birth), and were prepared by removing skin, organs, and brown fat. Then, they were dehydrated and fixed in 95 % ethanol for four days. To further remove fatty tissue and tissue permeabilization, specimens were exposed to acetone for one day. Consecutively, samples were transferred to Alcian blue and Alizarin red staining solutions. Later, they were exposed to potassium hydroxide (KOH) for three days, leading to tissue transparency. Finally, they were preserved in Glycerol 87%. The procedure can be adjusted depending on the size/age of the specimens (Rigueur and Lyons 2014).

### Dissection of branchial arches during development, RNA extraction, and RT-Digital Droplet^TM^ PCR (RT-ddPCR)

To study the level of expression of different genes triplicated on the DS mouse models, we performed RT-ddPCR (BioRad, Hercules, USA), a digital PCR used for absolute quantification that allows the partitioning of the cDNA samples obtained from the RT procedure up to 20,000 droplets of water-oil emulsions in which the amplification was performed (Lindner, Cayrou, Jacquot, et al. 2021).

For this, we collected the embryos of four pregnant females, Dp(16)1yey at E11.5 and four pregnant females of Dp(RNO11) (DS rat model with a complete duplication of chromosome 11) at E12.5. We obtained 20 embryos for each line (10 WT vs. 10 DS model, n=40) and dissected the frontonasal process, maxillary process, mandibular process, lateral and medial nasal process, and first pharyngeal arch. The dissected tissues are placed in cryogenic storage vials and quickly transferred to liquid nitrogen to avoid RNA decomposition. RNA extraction was performed using the RNeasy Plus Mini Kit (QIAGEN, Hilden, Germany), and RNA quality and concentration were assessed using Nanodrop (Thermo Fisher Scientific, Illkirh, France). cDNA synthesis was performed using the SuperScript™ VILO™ cDNA synthesis kit (Invitrogen), the final reaction is diluted five times and stored at −20 °C until use.

For the PCR reaction, 2 µL of the diluted cDNA samples are supplemented with 10 µL of Supermix 2X ddPCR (without dUTP, Bio-Rad, #1863024), 1 μl of target probe (ZEN™ FAM)/primers mix (final concentration of 750 nM of each primer and 250 nM of probe) and 1 μl of reference probe (ZEN™ HEX)/primer mix (final concentration of 750 nM of each primer and 250 nM of probe) obtaining a total volume of 20 μl. Once prepared, the samples are fractionated into droplets using the QX200 droplet generator (Bio-Rad). The PCR reaction can then be performed by transferring 40 µL of the samples to a 96-well plate. The fluorescence intensity of each droplet is then measured with the QX200 reader (Bio-Rad). Data are analyzed using QuantasoftTM Analysis Pro software (Version 1.0.596). More detailed protocol can be found in Lindner et al. 2021 (Lindner, Cayrou, Jacquot, et al. 2021).

Primers marked with specific fluorescent probes were designed using the PrimerQuest online web interface from IDT (https://www.idtdna.com/Primerquest/Home/Index) to target the genes of interest plus the housekeeping gene. Primers are blasted on the target gene map to verify that they span the exon/exon boundaries on the RNA. For mice and rats, the target genes were *Ripply3, Tbx1*, and *Dyrk1a*. The housekeeping gene for mice was *Tbp,* and for the rats, *Hprt1*. The primers are described below.

The primers used for mice were for *Ripply3* (ENSMUSG00000022941.9), forward AACGTCCGTGTGAGTCTTG, Reverse CTTTACTTACCCGTTTCAAAGCG, Probe ACACACATCGGGATCAAAGGGAGC (HEX); *Tbx1* (ENSMUSG00000009097.11), forward CTGTGGGACGAGTTCAATCA, reverse ACTACATGCTGCTCATGGAC, probe TCACCAAGGCAGGCAGACGAAT (FAM); *Dyrk1a* (ENSMUSG00000022897.16), forward GCAACTGCTCCTCTGAGAAA, reverse AACCTTCCGCTCCTTCTTATG, probe AAGAAGCGAAGACACCAACAGGGC (HEX); and housekeeping gene *Tbp* (ENSG00000112592.15), forward AAGAAAGGGAGAATCATGGACC, reverse GAGTAAGTCCTGTGCCGTAAG, probe CCTGAGCATAAGGTGGAAGGCT (FAM/HEX).

For the rat samples, primers were for *Ripply3* (ENSRNOG00000001684.7), forward GCTGATCTGACCAGAACTGAA, Reverse CGCTTTGAAATGGGCAAGTAA, Probe TTGGGAGGACCAACAAACCTTGGG (HEX); *Tbx1* (ENSRNOG00000001892.6), forward CAGTGGATGAAGCAGATCGTAT, reverse GGTATCTGTGCATGGAGTTAAGA, probe TCGTCCAGCAGGTTATTGGTGAGC (FAM); *Dyrk1a* (ENSRNOG00000001662.8), forward ACAGTTCCCATCATCACCAC, reverse TCCTGGGTAGAGGAGCTATTT, probe AATTGTAGACCCTTGGCCTGGTCC (HEX); and housekeeping gene *Hprt1* (ENSRNOG00000031367), forward TTTCCTTGGTCAAGCAGTACA, reverse TGGCCTGTATCCAACACTTC, probe ACCAGCAAGCTTGCAACCTTAACC (FAM/HEX).

### EdU Labeling

Depending on their capacity to proliferate, cells in the organism can be divided into three categories: proliferating cells, non-proliferating cells that have left the cell cycle, and quiescent cells capable of entering the cell cycle if necessary (Leblond 1964). Proliferating cells continue to progress through the different phases of the cell cycle (G1-> S-> G2->M). Daughter cells from a previous division immediately enter the next cell cycle. There are several specific markers for each phase of the cell cycle. For example, Phosphohistone 3 (PH3) corresponds to phosphorylated histone 3 and is found in mitotic cells. EdU is a thymidine analog that can be incorporated into DNA during replication (S phase of the cell cycle). We can define the percentage of proliferating cells and cell cycle progression using these two markers.

First, pregnant females Dp(16)1Yey at E8.5 stage were injected intraperitoneally with EdU (SIGMA, ref. E9386; diluted in NaCl 0.9%, final concentration 7.5µg/µL; volume injected: 41ug for each milligram of animal weight). Then, embryos were collected 24 hours after EdU injection (E9.5). After, embryos were fixed with 4% paraformaldehyde and embedded in Shandon Cryomatrix Frozen Embedding Medium (Thermo Scientific™). Frozen sagittal sections (14 μm) were cut using a Leica CM3050 S cryostat and placed on Superfrost Plus™ slides for immunohistochemistry.

The immunohistochemistry for EdU was performed following the protocol described in the Kit de cellular proliferation EdU Click-iT™ for imaging, Alexa Fluor™ 555 (REF: C10338). For PH3 we used as primary antibody the Anti-phospho-Histone H3 (Ser10) Antibody, Mitosis Marker (Merck Millipore, REF: 06-570, 1:500) and secondary antibody the Donkey anti-Rabbit IgG (H+L), Alexa Fluor™ 647 (Invitrogen, REF: A-31573, 1:500). For the nuclear marker we used Hoechst nucleic acid stain (Invitrogen, REF: H3570, 1:2000). Quantitative analyses comparing wild-type and mutant embryos. The percentage of EdU-positive cells determined the proliferation index (the number of proliferating cells relative to the total number of cells, labeled with Hoechst) in the area of interest. In addition, the ratio of EdU-positive/PH3-positive cells allow to assess how many proliferating cells have progressed from S phase to G2/M phase (mitotic cells), thus defining the mitotic index.

## RESULTS

### Contribution of *Lipi-Zbtb21* region to DS craniofacial features in Dp(16)1Yey mouse model

In individuals with DS, sexual dimorphism is observed since early age, with higher measurements in males than in females, but the growth rate remains unchanged. However, in the studies where cephalometric superimposition variables were analyzed, these differences did not appear. This might be because of the low magnitude of the superimposition measurements; making it difficult to determine significant differences (Alio, Lorenzo, and Iglesias 2008; Vicente et al. 2020).

Previous studies in mice have shown no sex differences in the shape of the cranium and only a subtle difference in the shape of the mandible (Toussaint et al. 2021). Importantly, for both the cranium and mandible, the effect of genotype was stronger than sex for Dp1Tyb mice and the other strains (Redhead et al. 2023). Considering this information, we decided to verify this observation and used both sexes together in the CF analysis of Dp(16)1Yey.

First we analyzed the Dp(16)1Yey DS mouse models to compare them with the other DS model Dp(16)1Tyb (Redhead et al. 2023). Then, we performed morphometric analysis of Dp(16)1Yey on adult samples. We observed significant changes in form and shape difference matrix that can be understood as an overall reduction of dimensions (microcephaly) and smaller mandible (Fig S1). For a more detailed investigation of the patterns of displacements of landmarks and their dimensionality, we used principal component analysis (PCA). In the skull and mandible, the PCA of Dp(16)1Yey showed significant differences in the dimensionality versus the WT group (Figure 1A).

**Figure 1.**
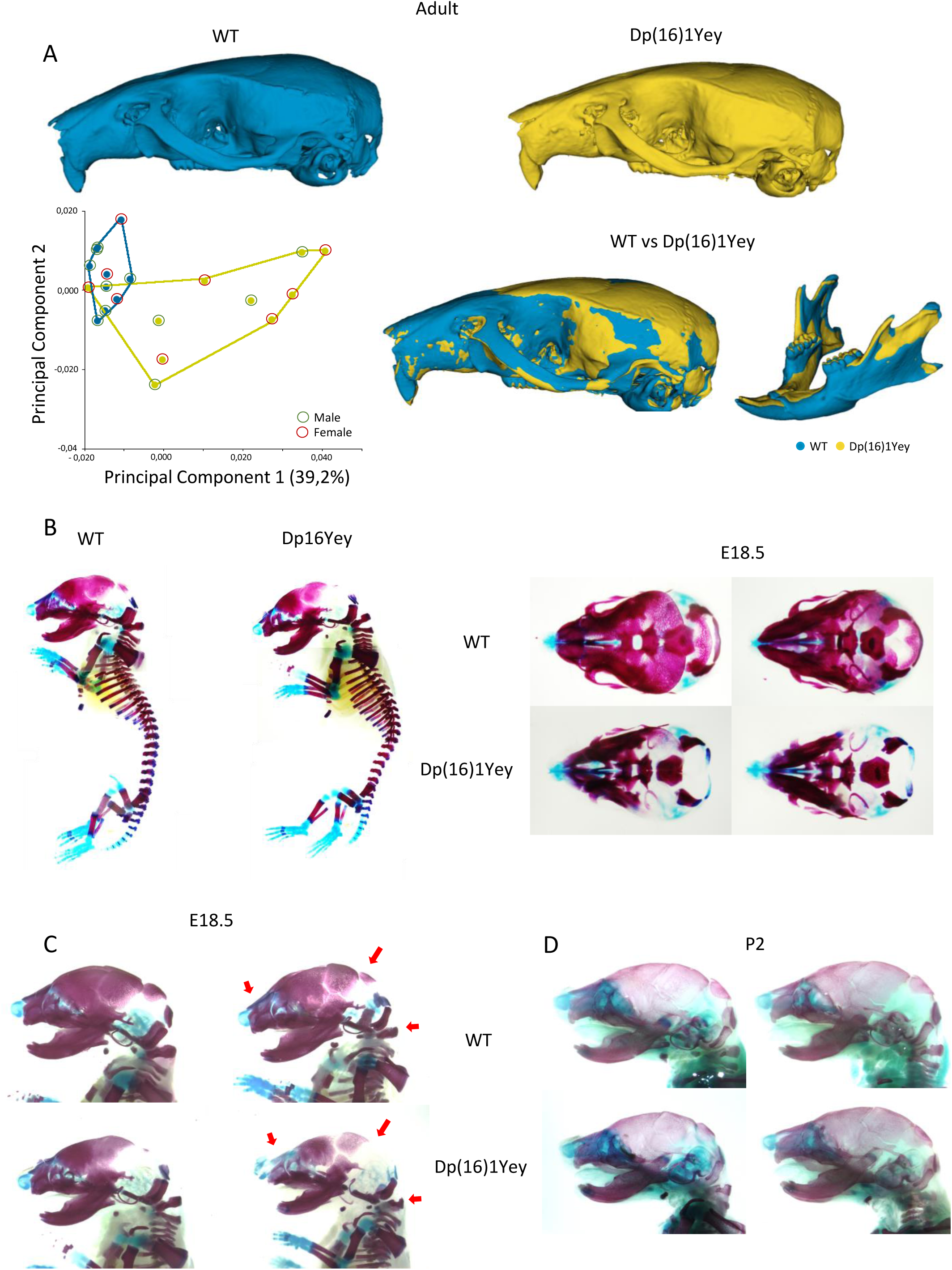
Alteration of DS craniofacial features in Dp(16)1Yey is observed during late development. **(A)** Morphometric analysis of the cranium and mandible of adult Dp(16)1Yey vs control. WT is in light blue and Dp(16)1Yey in yellow. PCA (first two components) of general Procrustes analysis of aligned cranium shapes, using data from females (red circle) and males (green circle) mice. The percentage of variance in PC1 corresponds to 39.2%. In the figure WT vs Dp(16)1Yey shape difference warping, with in blue the bones with decreased dimensions in Dp(16)1Yey (midface hypoplasia and mandible), and in yellow, the bones with increased dimensions (neurocranium). **(B-D)** Skeletal staining with alizarin red and alcian blue. Comparison WT vs Dp(16)1yey E18.5 whole body, upper (top) and lower (bottom) magnification of the skull showing less mineralization in temporal, parietal, intraparietal, and occipital bones. **(C)** Magnification of lateral view of skeletal staining, WT vs Dp(16)1Yey at E18.5 (2 individuals). Red arrows showing less mineralization in nasal bones, neurocranium, and form defect in atlas vertebrae. **(D)** Lateral view of skeletal staining of two WT vs Dp(16)1Yey individuals showing normal ossification pattern at P2.

To track the landmarks associated with a significant change and understand where they are located in the CF structures, “EDMA FORM or SHAPE Influence landmark analysis” was performed (Cole et Richtsmeier 1998). The purpose of this test is to search which landmarks present a Relative Euclidean distance (RED) > 1.05 or < 0.95 (outside of the confidence interval 97,8%), meaning which landmarks show a bigger difference in linear distances between every landmark and in what direction. On one side, the most influential landmarks with decreased dimensions corresponded to the ones from the maxillary bones, mandible, premaxilla, frontal, temporal (with the squamosal portion), and occipital bone. On the other side, landmarks with a significant increase of dimensions were in the cranial vault, the parietal bone, and the intraparietal bone (Fig. S1).

For a more detailed investigation of the patterns of displacements of landmarks and their dimensionality, we used PCA. In skull and mandible, the PCA of Dp(16)1Yey showed significant differences in the dimensionality versus the WT group (Figure 1A). Also, using Procrustes average models of the different populations to perform a voxel analysis, we found that the key aspects of the Dp(16)1Yey phenotype correspond to a decrease in the dimensions of the midface, defining a midface hypoplasia and a short nasal region. In the neurocranium, an increase of dimensions in the lateral width was found, with a reduction in the occipital bone, leading to a shortening of the anteroposterior axis (brachycephaly). In the case of the mandible, we found a decrease in the width of the ramus, body, incisor alveolus, and molar alveolus and increased lateral dimension in coronoid and condylar process (expected by the skull brachycephaly; Figure 1A).

Besides, individuals with DS also experience a low bone mass associated with reduced osteoblast activity and bone turnover (McKelvey et al. 2013). Knowing this, plus the information obtained in the craniofacial analysis, we proposed that these significant changes could affect bone ossification during development. To address this, we performed a skeletal/cartilage staining with alizarin red and alcian blue in Dp(16)1Yey in embryonic stage (E) 18.5 and P2. At E18.5, mutant embryos exhibit a defect in the mineralization in the parietal bones, intraparietal, nasal bone, and atlas compared to WT (Figure 1B-C). Interestingly, no more phenotype was observed at P2 (Figure 1D). Altogether, the origin of the CF morphological changes in the Dp(16)1Yey are similar to Dp(16)1Tyb (Toussaint et al. 2021) and probably originate during pre-natal development in the mouse.

### Dissection of CF phenotype: Mapping the location of dosage-sensitive genes inside *Lipi-Zbtb21,* that cause the craniofacial dysmorphology using a new panel of mouse models

To elucidate the location of dosage-sensitive genes that are predominantly involved in the craniofacial dysmorphology of Dp(16)1Yey mice, we took advantage of a new panel of 7 mouse models with shorter segmental duplications covering the Mmu16: Dp(16)7Yah, Dp(16)8Yah, Dp(16)9Yah, Dp(16)10Yah, Dp(16)11Yah, Dp(16)12Yah and Dp(16)13Yah. We analyzed their CF morphology and compared duplication versus their control wild-type (WT) littermates (Fig. 2A).

**Figure 2.**
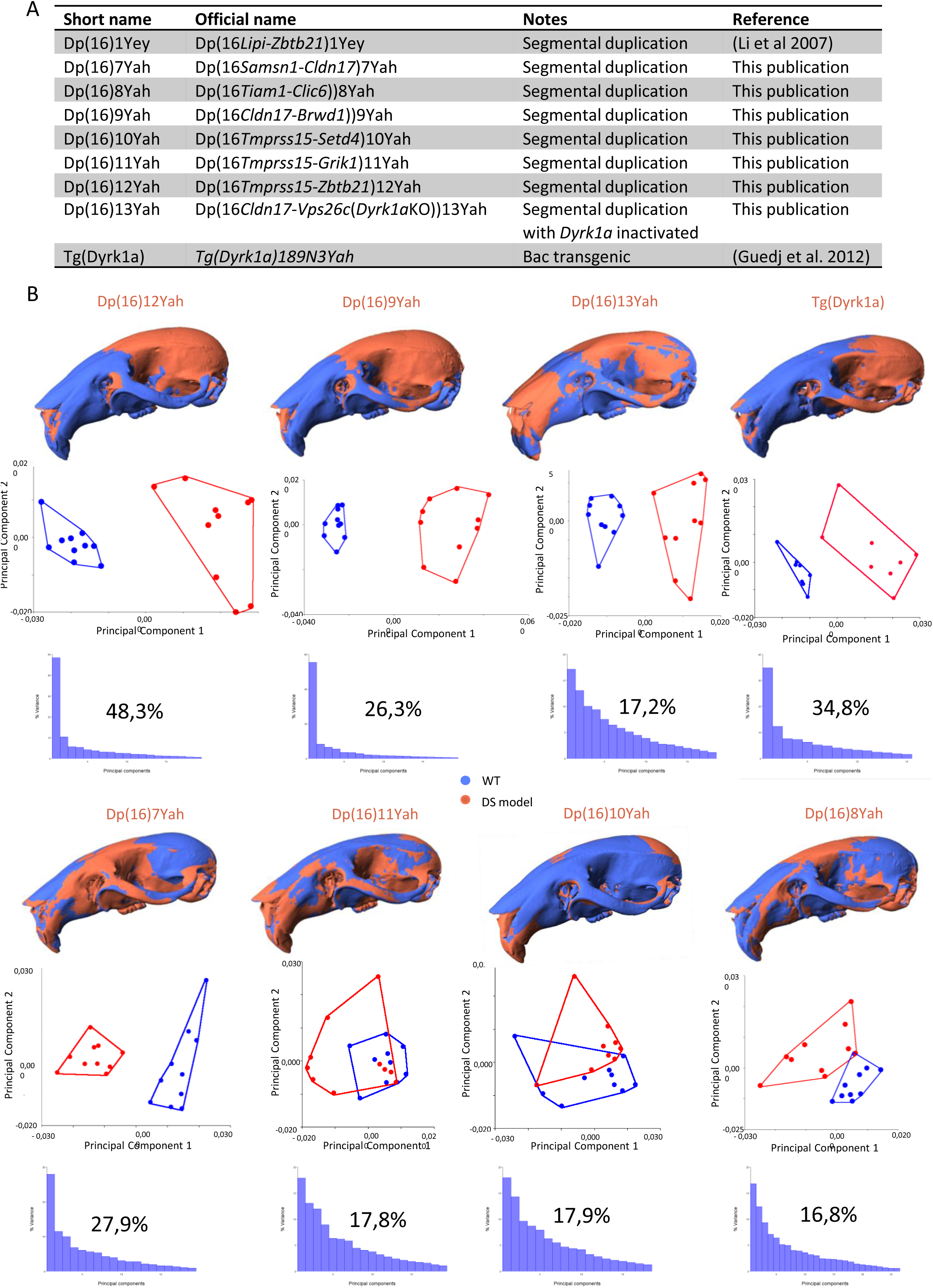
New DS mouse model panel to identify genetic regions whose overdosage causes the craniofacial dysmorphology of Dp(16)1Yey. **(A)** Summary table of a new panel of DS mouse model and their segmental duplication of the genetic interval. **(B)** Morphometric analysis of the cranium of the new panel of DS mouse models, plus Tg(Dyrk1a). Shape difference warping highlighted, in blue, the bones with decreased dimensions in DS models, and in red, the bones with increased dimensions. PCA (first two components) of general Procrustes analysis of aligned cranium shapes for every model with the percentage of variance graphic in PC1.

Performing the identical craniometric analysis as previously described, we found significant skull changes in form and shape in all the models except Dp(16)8Yah (Fig. S2). Multivariable analysis using PCAs showed changes in the same direction along principal component 1 as seen in Dp(16)1Yey mice (Fig. 2B). A major contribution of the principal component 1 (PC1) for Dp(16)12Yah, Tg(*Dyrk1a)*, was found as in Dp(16)1Yey, then Dp(16)7Yah and Dp(16)9Yah.

Similar changes in shape and form in the skull, with an overall reduction in midface region and a strong brachycephaly, were observed in Dp(16)1Yey, Dp(16)9Yah and Dp(16)12Yah (Fig. 2B). The Dp(16)13Yah model also presented a reduction in the midface region. Still, the premaxilla and nasal region were not reduced, and the brachycephaly was not seen (Fig 2B). The Tg(*Dyrk1a)* model, overexpressing *Dyrk1a* alone, showed strong brachycephaly and reduced premaxilla and nasal bone.

In contrast, we found an inverse phenotype in Dp(16)7Yah compared to Dp(16)1Yey. We scored increased dimensions in the midface and decreased dimensions in the neurocranium; similar changes were found in Dp(16)11Yah but were less significant. Dp(16)10Yah presented an overall reduction of head dimensions, partially observed in the Dp(16)8Yah with a larger midface part. Still, an increase in premaxilla and occipital bone size, resulting in an elongation of the anteroposterior axis in Dp(16)10Yah also observed in Dp(16)13Yah (Fig. 2B). The PCAs showed changes in the inversion direction along principal component 1 as seen in Dp(16)1Yey mice (Fig. 2B).

For the mandibles, as in the skull, significant changes in form and shape were found in all the lines (Fig S3). Here, the changes were different, although the most important changes were found in the same DS models, namely Dp(16)9Yah, Dp(16)13Yah, Dp(16)12Yah, and Dp(16)7Yah. In Dp(16)9Yah and Dp(16)12Yah, a reduction in the lateral width on the body of the mandible, ramus, molar and incisor alveoli, but an increase in the dimension in the condylar process (coincident with the brachycephaly found in the neurocranium) were detected; these changes were similar to the ones found in Dp(16)1Yey (Fig S3). In the case of Dp(16)13Yah, this model also presented a similar phenotype but with increased dimensions in the ramus (apart from the condylar process) (Fig. S3). Dp(16)7Yah had increased dimensions in the body of the mandible, ramus, molar alveolus, and condylar process but maintained the reduced dimensions of the incisor alveolus. A similar phenotype is found in Dp(16)11Yah, but dimensions decrease in the angular process. Also, in Dp(16)10Yah, significant changes were found in the condylar process, presenting a reduction in the lateral width (Fig. 2B).

The analysis of this new mouse panel allows us 1) to dissect the contribution of several Mmu16 region overdosages to CF phenotypes in DS and 2) to show the differences in their contribution to the cranium and mandible CF phenotypes.

### Candidate genes for CF DS phenotype in DS rodent models

We focused on a new chromosomal region selected from the previous morphometric analysis, encompassing 15 genes between *Setd4 –Dyrk1a* genes (Fig 3A). Inside this region, based on scientific literature and a data set from the FaceBase Consortium (Hong Li 2017), we isolated a set of candidate genes. Among these genes, *Dyrk1a* (Dual Specificity Tyrosine Phosphorylation Regulated Kinase 1A) and *Ripply3* (Ripply Transcriptional Repressor 3) seemed promising targets for CF in DS.

**Figure 3.**
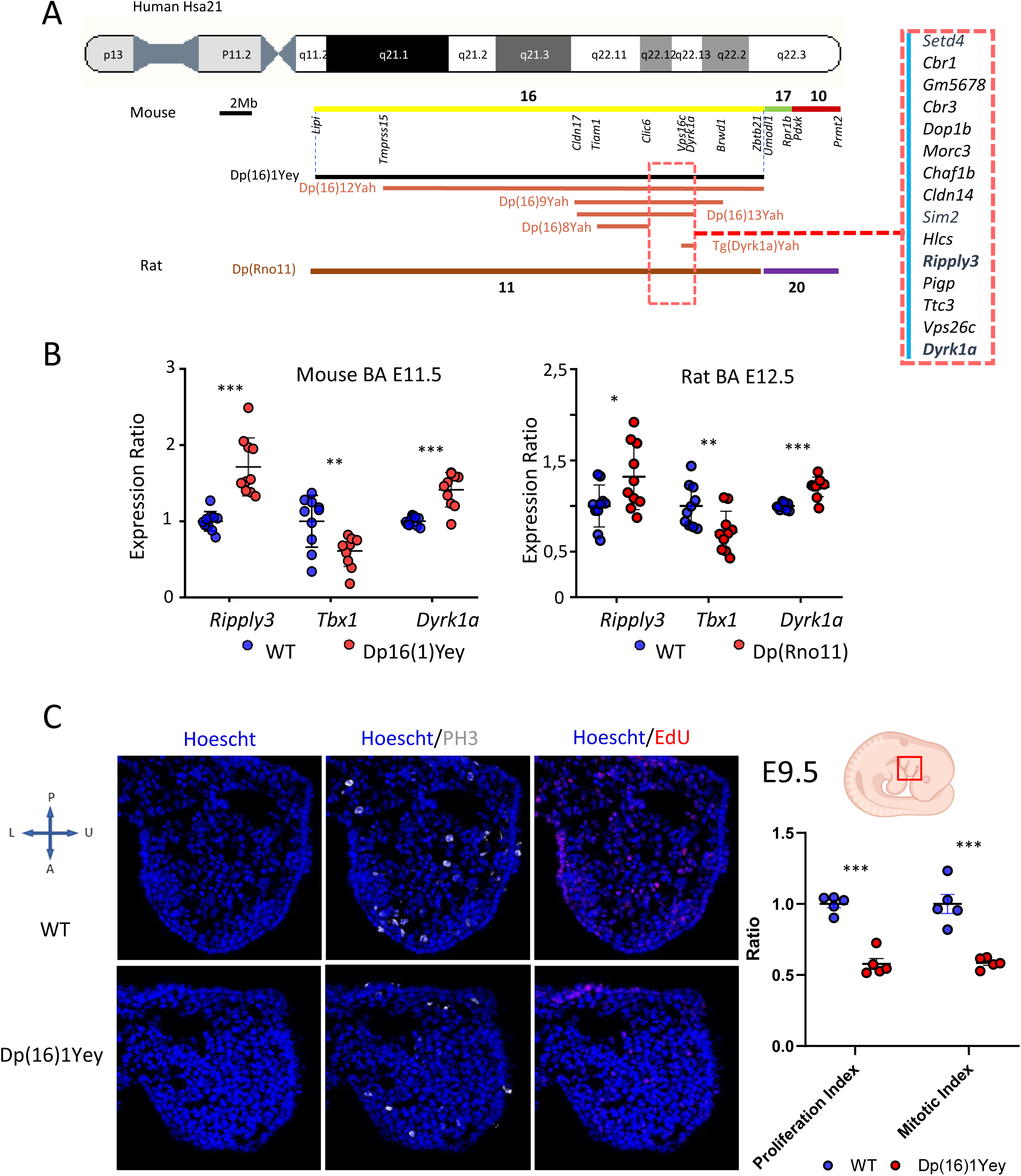
Genetic mapping identifies a new chromosomic region and dosage-sensitive genes for the DS craniofacial phenotype. **(A)** Scheme of the relative position of the new mouse and rat models in HSA21, and the new chromosomic region of interest for DS CF phenotype (red square). **(B)** RT-ddPCR results graphics for Mouse model Dp(16)1Yey E11.5 in the left, the gene expression ratio shows statistically significant overexpression of the triplicated *Dyrk1a* (***, p<0.000) and *Ripply3* (***, p<0.000) and a downregulation of *Tbx1* (**, p<0.005). In the right, the graph for the Rat model Dp(RNO11) E12.5. The gene expression ratio shows statistically significant overexpression of the triplicated *Dyrk1a* (***, p<0.000) and *Ripply3* (*, p< 0,02) and a downregulation of *Tbx1* (**, p<0.01). **(C)** Edu, PH3, and Hoechst unrevealed defects in the proliferation and mitosis of the NCC derivates in the 1st branchial arch during craniofacial development in Dp(16)1Yey. In the right graphic, the proliferation (***, p<0.000) and mitotic (***, p<0.000) indexes were significantly reduced in Dp(16)1Yey.

To explore this hypothesis, we analyzed the expression levels of *Dyrk1a, Ripply3*, and *Tbx1* via Droplet digital polymerase chain reaction (RT-ddPCR) in Dp(16)1Yey branchial arches and frontal process at E11.5. The stage is just after the neural crest-derived mesenchymal cells are differentiated into different bones in the facial region (Everson et al. 2018). In the Dp(16)1Yey samples, an overexpression of *Dyrk1a* and *Ripply3* is detected in the craniofacial precursor tissues in mutant versus wild-type. Concomitantly, *Tbx1* was downregulated in trisomic model versus control.

We also did similar expression studies in the Dp(RNO11) rat model with complete duplication of *Lipi-Zbtb21* region in the rat chromosome 11 (Birling et al. 2017) at embryonic stage E12.5 (homologous to the mouse stage). Similar results were found in the rat DS models with overexpression of *Dyrk1a* and *Ripply3* and down-regulation of *Tbx1* (Figure 4B). Overall, the *Ripply3* overexpression due to the triplication of the gene has the same consequence with a reduced expression of *Tbx1* in the branchial arches.

**Figure 4.**
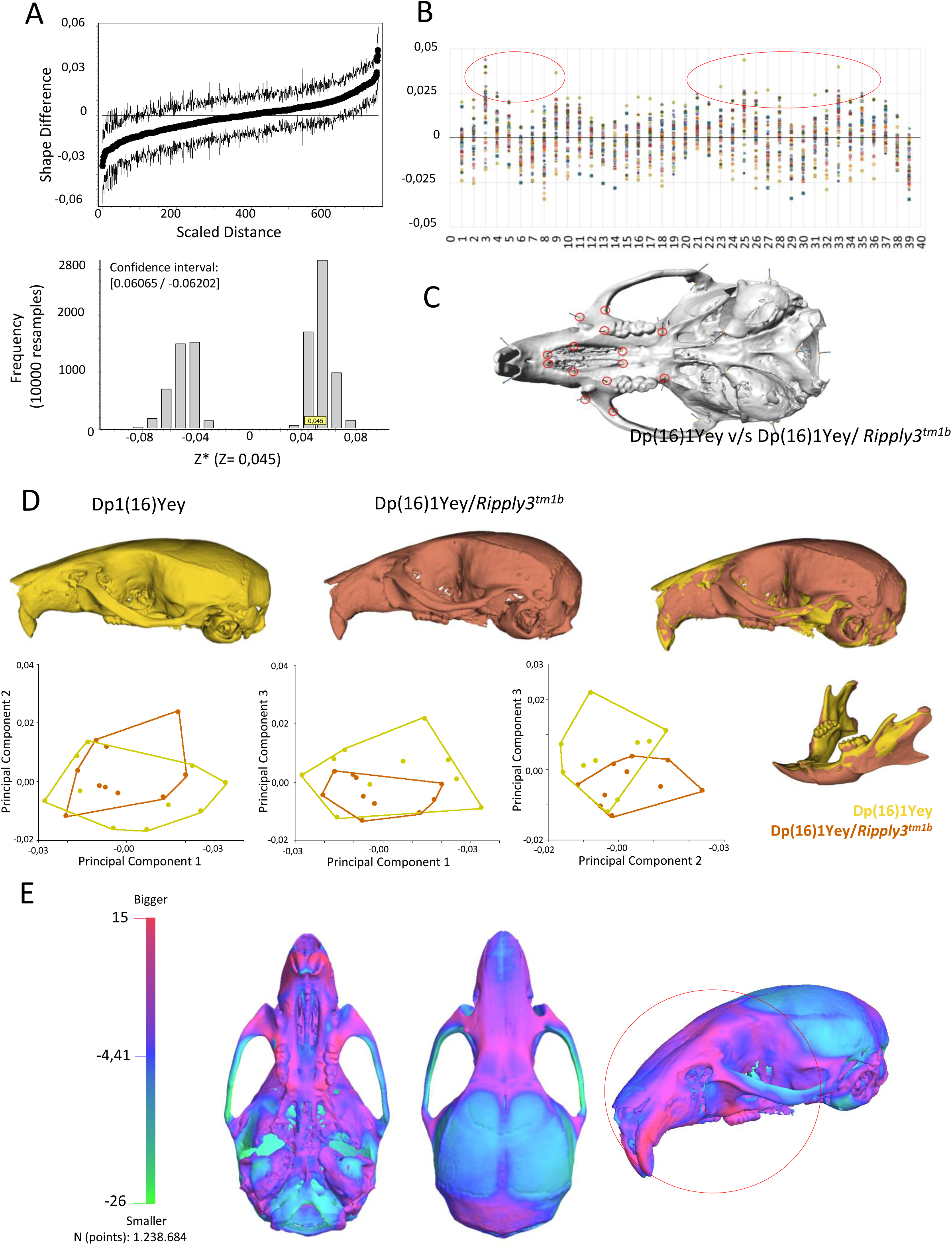
Shape rescue in the midface phenotype in the Dp(16)1Yey/ *Ripply3^tm1b^* compound mutant. **(A)** Shape difference matrix based on Dp(16)1Yey vs Dp(16)1Yey/ *Ripply3^tm1b^* comparison. Confidence interval and Frequency Bootstrap graphics with 10.000 resamplings showed significant shape changes (Z statistics = 0,045, CI = [0.06065 / - 0.06202]). Influence landmarks analysis to show the most influential landmarks (red circle) that lead to significant changes in Dp(16)1Yey/ Ripply3^tm1b^ vs Dp(16)1Yey in **(B)** with their position **(C)** located in the midface. **(D)** Shape difference warping of Dp(16)1Yey vs Dp(16)1Yey/ *Ripply3^tm1b^*. Dp(16)1yey is displayed in yellow and Dp(16)1Yey/ *Ripply3^tm1b^* in orange. The bones with increased dimensions are in orange in the cranium figure of Dp(16)1Yey vs Dp(16)1Yey/*Ripply3^tm1b^* warping. Maxillary bone, premaxilla, alveolar process and neurocranium are similar to that found in Dp(16)1Yey). In yellow, the bones with decreased dimensions (nasal bone and skull base). In the mandible of Dp(16)1Yey vs Dp(16)1Yey/ *Ripply3^tm1b^*warping, the mandible presented the same changes found in Dp(16)1yey. PCA (first three components) of general Procrustes analysis of aligned cranium shapes. Showing significant changes in PC1/PC2, PC3/PC1 and PC3/PC2. **(E)** Comparison of Dp(16)1Yey average model vs Dp(16)1Yey/Ripply3*^tm1b^* average model with a 3D heatmap from 3dMD Vultus ® software analysis. Pink/red shows increased shape dimensions in all the structures corresponding to the midface. In light blue, the structures with no significant changes. On the left, the histogram of every point distance evaluated (in this case, more than 1.238.684 points) and the surface differences with the color code for the increase-decrease dimensions (Red to green).

### Role of *Dyrk1a* overdosage in the increased dimensions in neurocranium (brachycephaly) on DS mouse models

*Dyrk1a* has been implicated in several DS phenotypes and craniofacial abnormalities (Guedj et al. 2012; McElyea et al. 2016; Redhead et al. 2023; Starbuck et al. 2021) and in the last decade has become one of the top candidate gene in DS for therapeutic intervention (De la Torre et al. 2014; Atas-Ozcan et al. 2021). Here we took advantage of Tg(*Dyrk1a*) - a model with 3 copies of *Dyrk1a* (Guedj et al. 2012) - and compared with the results of the new model Dp(16)13Yah where Dyrk1a is not overexpressed, to confirm the role of *Dyrk1a* in the development of the DS CF phenotype. Brachycephaly and higher dimensions in the neurocranium were present in Tg(*Dyrk1a*), but also a reduction in the midface region, a phenotype very similar to the one observed in Dp(16)1Yey (Fig. 2). In Dp(16)13Yah, a reduction in the midface region was also seen but focussed in the maxillary bones, the premaxilla and nasal region were not reduced and the strong brachycephaly was absent. Overall, this comparison confirmed the hypothesis and the role of *Dyrk1a* overdosage in inducing the brachycephaly.

### *Dyrk1a* and *Ripply3* overdosages affect the proliferation and mitosis of the NCC derivates in the first branchial arch during craniofacial development

In Tg(Dyrk1a) and in the new panel of mouse models (specifically in Dp(16)9Yah, Dp(16)12Yah and Dp(16)13Yah), significant changes were observed in the midface region, in structures that share the same embryonic origin, the neural crest cells (NCC) (Richtsmeier and Flaherty 2013). These findings, considering the relation of *Dyrk1a* with an altered expression of critical craniofacial regulators fundamental for cranial neural crest development (Johnson et al. 2024) allows us to postulate that *Dyrk1a* and probably *Ripply3* overdosages could affect the proliferation of the NCC derivates during craniofacial development.

To demonstrate this, we monitored the proliferation of NCC with 5-Ethynyl-2’-deoxyuridine (EdU) a thymidine analogue incorporated into the DNA during replication (Tucker et al. 2010). Compared to controls, immunohistological analysis was used to detect the proliferation of NCC derivates in the first branchial arch of the Dp(16)1Yey embryos. In addition, quantitative analyses defined the proliferation and mitotic index (Harris et al. 2018). A reduced proliferation and mitotic index in the first branchial arch were detected (Fig. 3C). Thus, overexpression of triplicated genes from Dp(16)1Yey, including *Dyrk1a* and *Ripply3*, lead to the reduced proliferation of mesenchyme cells from the branchial arches.

### Confirmed role of *Ripply3* overdosage during midface development in DS mouse models

To confirm the role of *Ripply3* in midface hypoplasia, we crossed mice with a loss-of-function allele (*Ripply3^tm1b/+^*) obtained from the IMPC initiative (www.mousephenotype.org) with the Dp(16)1Yey line. We obtained Dp(16)1Yey/*Ripply3^tm1b^* males carrying a trisomy of all the genes present in Mmu16 but with only two functional copies of *Ripply3*, this to rescue the midface hypoplasia of Dp(16)1Yey. Morphometric analysis of Dp(16)1Yey/*Ripply3^tm1b^* showed significant shape changes in the skull and mandibles compared to Dp(16)1Yey. In the SDM influence landmarks analysis, we can observe that the most influence landmarks leading to increased dimensions are located in the midface (Fig. 4C). The skull voxel analysis showed the expected result with a similar change in the neurocranium to that present in Dp(16)1Yey. We noticed a phenotypical shape rescue in the midface, with increased dimension in maxillary bones, alveolar process, and premaxilla. The mandible presented the same changes found in Dp(16)1yey (Fig. 4D). To confirm this result and obtain further details, we mapped the surface differences in 3dMD Vultus® software. Using the average model of each population, Dp(16)1Yey average model vs Dp(16)1Yey/*Ripply3^tm1b^* average model, we performed a superimposition of the 3D models employing landmarks to have an exact matching (Fig 4E). Once the comparison was made, we obtained a histogram of every distance evaluated (in this case more than 1.238.684 points) and a 3D heatmap with the surface differences. In the heat map, we could observe the increased dimensions in all the bones corresponding to the red regions’ midface. In light blue, the structures that did not present significant changes (Fig. 4E).

### Integrative multivariate analysis of the craniofacial studies unravels four different subgroups of DS models

So far, after doing separate analyses of the DS models with their mutant and control littermates, we decided to combine all the data and see whether we could discriminate different contributions. As shown in Fig. 5, all the wild-type controls clustered together for the cranium and the mandible, with a higher variation, probably due to the altered position of some landmarks.

**Figure 5.**
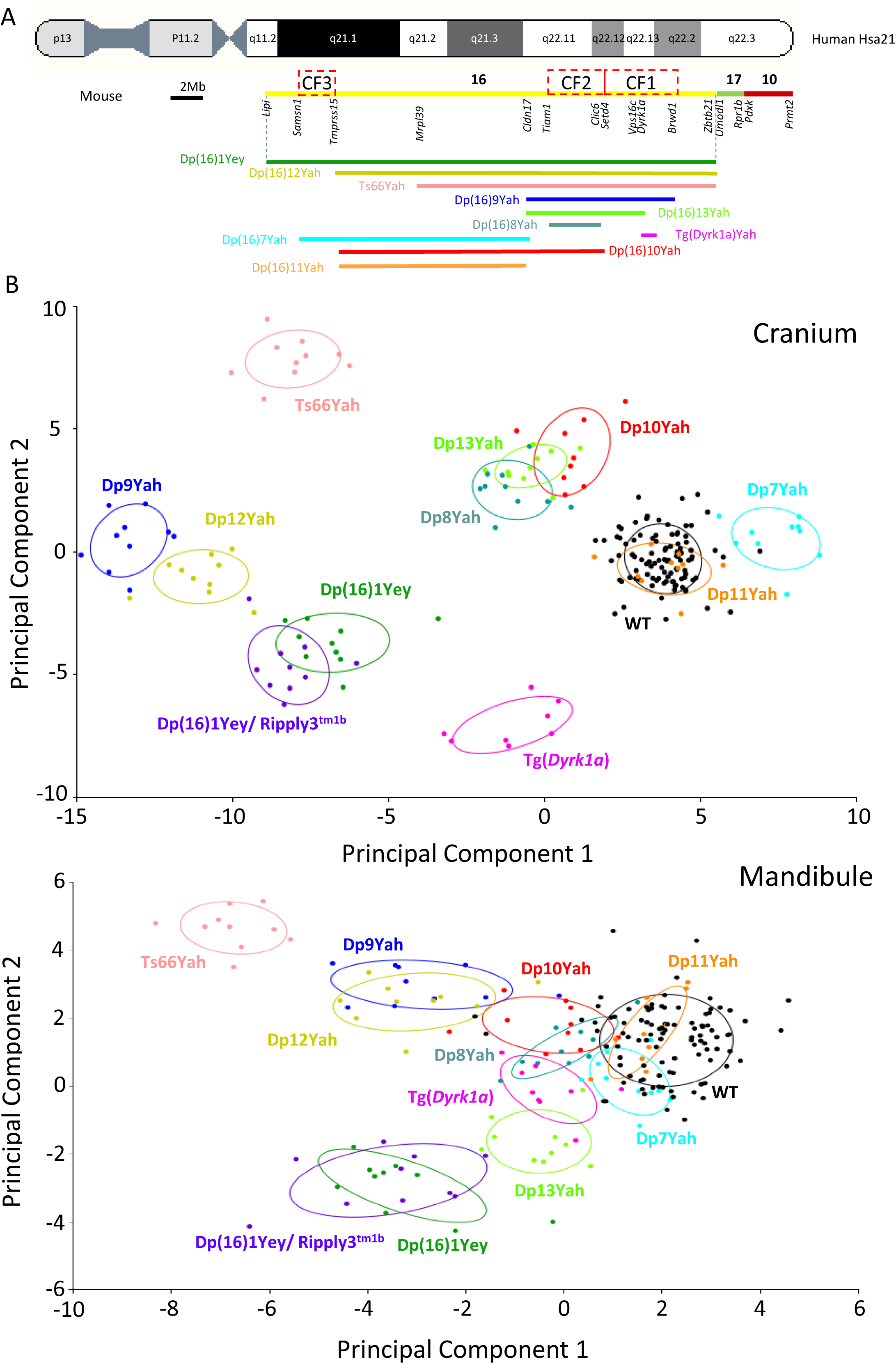
Integrative multivariate analysis of the craniofacial studies. **(A)** Schematic representation of DS models and their relative position to HSA21, showing the 3 new CF regions (red squares) involved in the DS CF phenotype located on Mmu16. **(B)** Integrative multivariate analysis of all the models used in this study, plus Ts66Yah vs wild-type. The PCAs correspond to a canonical variate analysis (Procrustes Distance Multiple Permutations tests at 1000 iterations). DS strains show significant differences compared with their wild-type controls. DS models were separated into four main groups with the cranium PCA graph, whereas for the mandible, the graph showed a prominent group of 5 models close to the wild-type and two branches separated on PC2.

Focusing on the skull, the DS models were separated into four main groups according to the first two dimensions. One is composed of the Dp(16)1Yey, Dp(16)1Yey/*Ripply3^tm1b^*, a second of Dp(16)12Yah and Dp(16)9Yah, a third with Dp(16)10Yah, Dp(16)13Yah and Dp(16)8Yah, and the Dp(16)11Yah mixed with the wild-type while the Ts66Yah, the Tg(Dyrk1a) are on the same side of the DS models and the Dp(16)7Yah stayed apart from the other on PC1. Somehow this distribution of models reflected the complexity of genetic interactions for the cranium phenotypes with at least 3 minimal Mmu16 regions: CF1 from *Setd4* to *Brwd1,* involving notably *Dyrk1a* and *Ripply3*, a second CF2 mapping to *Tiam1-Clic6*, and a third CF3 *Samsn1-Tmprss15* responsible for the Dp(16)7Yah phenotypes.

For the mandible, the graph showed another complexity with a main group of 5 models closed to the wild-type (Dp(16)11Yah, Dp(16)7Yah, Dp(16)10Yah, Dp(16)8Yah, Tg(Dyrk1a)) then two branches separated on PC2 with on one side the Dp(16)1Yey, Dp(16)1Yey/*Ripply3^tm1b^*, linked to the wt-like group with Dp(16)13Yah, and on the other side Dp(16)12Yah and Dp(16)9Yah. The Ts66Yah being kept alone. Interestingly the Dp(16)1Yey, Dp(16)1Yey/*Ripply3^tm1b^*, Dp(16)12Yah and Dp(16)9Yah, are closer for cranium changes, but they are well-separated for the mandibular phenotypes. These differences could come from various interacting regions contributing to the cranium and mandibular phenotypes.

## DISCUSSION

In DS individuals, craniofacial dysmorphism is almost 100% penetrant, but the contributive genetic and developmental factors were unclear. The DS CF phenotype typically encompasses microcephaly, a small midface, a reduced mediolateral orbital region, reduced bizygomatic breadth, a small maxilla, brachycephaly (a relatively wide neurocranium), and a small mandible (Olson et al. 2004).

We used Dp(16)1Yey to study this characteristic phenotype. This model carries a complete duplication of Mmu16 (Li et al. 2007) and a previously well-described DS-like craniofacial phenotype (Starbuck et al. 2014). Our results reproduce the same findings using a standard craniofacial analysis plus a new voxel analysis, where we could observe the principal changes in the skull and mandibles in 3D. We found that the changes were correlated with the human DS CF phenotype, making Dp(16)1Yey the appropriate model to study DS CF phenotype. Besides, we performed skeletal staining where we could identify a defect in intramembranous ossification, the results obtained at different embryonic stages allowed us to postulate that the changes observed at E18.5 did correspond to a delay and not to a continuous defect in intramembranous ossification. This defect could be related to a problem in the differentiation of mesenchymal cells into osteoblasts or in osteoblast proliferation with a subsequent attenuated osteoblast function (Thomas et al. 2021).

Here we demonstrated that the craniofacial dysmorphism found in Dp(16)1Yey, correlated with the human DS CF phenotype, and we mapped three distinct chromosomic regions of Mmu16. Using a new panel of DS mouse models with specific CF phenotypes (Table 1), we could now identify new dosage-sensitive regions and genes responsible for DS CF phenotype. Saying this, the Ts66Yah behaved independently according to dosage effects in CF phenotypes compared to lines with segmental duplication. This may reflect a unique contribution of the minichromosome on the CF severity while the gene content is close to Dp(16)9Yah. Further experiments with new models will help to confirm the role of the segregating chromosome in CF DS phenotypes (Xing et al. 2023).

**Table 1:**
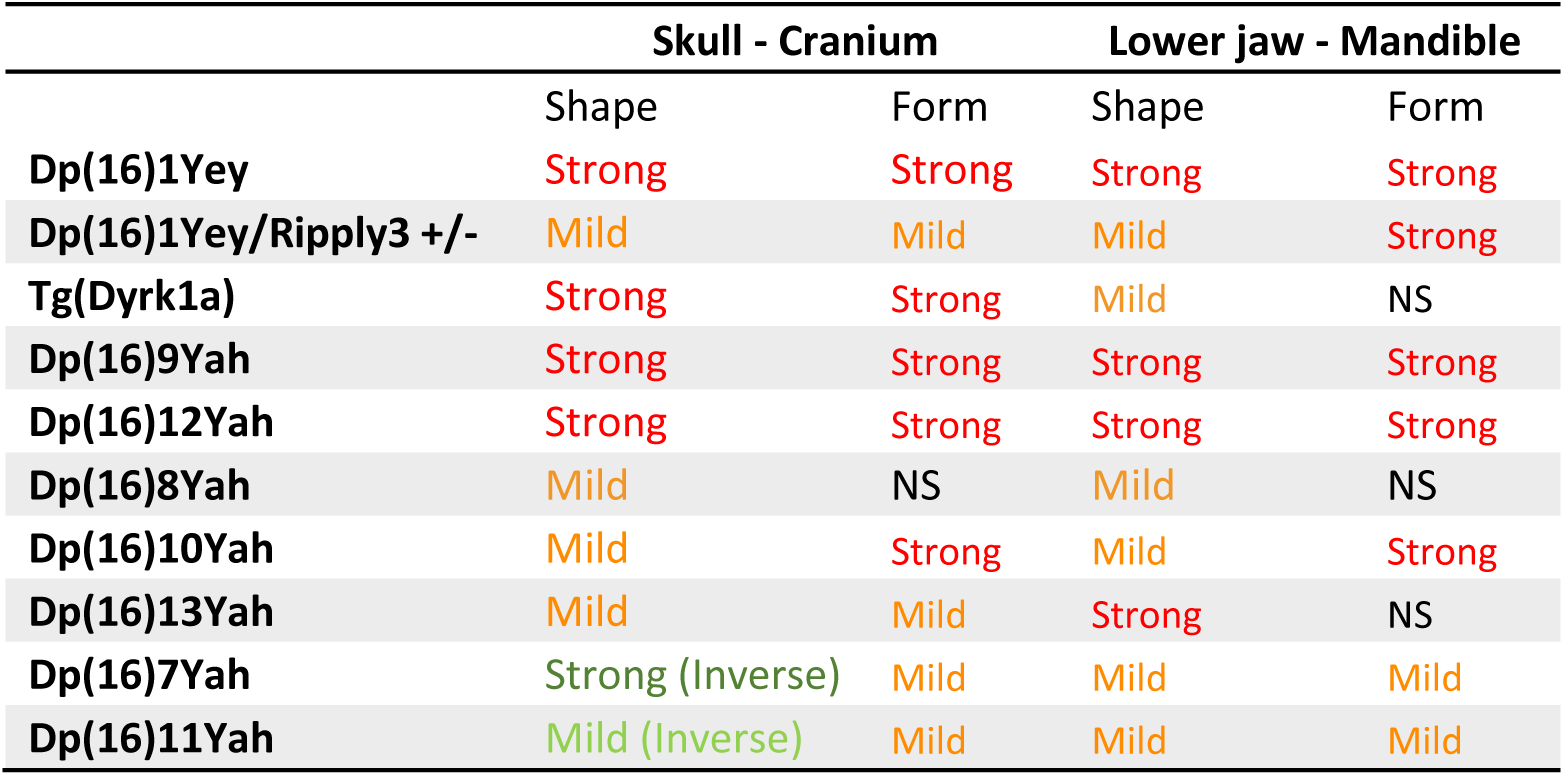
Summary of the CF phenotypes observed in DS models.

This report excludes the *Tmprss15-Grik1* regions, triplicated in Dp(16)11Yah alone, with almost no effect on CF form and shape. This agrees with the Dp(16)9Tyb lack of CF phenotype (Redhead et al. 2023). On one hand, two of the three CF regions defined here, CF1 and CF3, settled the telomeric regions described by Redhead et al. (2023). Nevertheless, we found the contribution of the most centromeric part CF3 involved in cranium enlargement as new. It could be slightly artificial as this effect was not seen in the Dp(16)1Yey, but could also be due to an effect specific to this region while not triplicated with CF2 and CF3. Only more detailed investigations with new models would allow us to discriminate the genetic interaction of the 3 CF regions. On the other hand, while the overdosage of *Dyrk1a* was crucial for the Dp(16)1Tyb phenotypes (Redhead et al. 2023), the sole overexpression of *Dyrk1a* in the Tg(Dyrk1a) line only replicated well the skull phenotype, more precisely the brachycephaly. Conversely, using PCA, transgenic individuals were closer to the WT group for the lower jaw. Recently, *Dyrk1a* has been identified as one of the genes required in three copies to cause CF dysmorphology in mouse models of DS, and the use of DYRK1A inhibitors or genetic knockout of DYRK1A has been shown to rescue the skull and jaw malformations (McElyea et al. 2016; Redhead et al. 2023). However, Johnson et al. in 2024, showed that a decrease in *Dyrk1a* in Xenopus resulted in craniofacial malformations, altered expression of critical craniofacial regulators as *Pax3* and *Sox9* fundamental for cranial neural crest development, and presented altered retinoic acid, hedgehog, nuclear factor of activated T cells (*NFAT*), *Notch* and *WNT* signaling pathways. These results indicate that DYRK1A function is critical for early craniofacial development and must properly regulate the expression of specific craniofacial regulators in the branchial arches (Johnson et al. 2024). We achieved to demonstrate, thanks to the analysis in Tg(*Dyrk1a*) and Dp(16)13Yah, that 3 copies of *Dyrk1a* are necessary to induce the brachycephaly found in DS. Thus, *Dyrk1a* overdosage is essential and sufficient for brachycephaly, but other genes are responsible for the mandibular phenotypes observed in DS.

Our other candidate gene*, Ripply3*, is a transcriptional corepressor, acting as a negative regulator of the transcriptional activity of *Tbx1* and playing a role in the development of the pharyngeal apparatus and derivatives (Okubo et al. 2011). *Tbx1* Is the first dosage-sensitive gene identified in the DiGeorge syndrome (DGS)/velocardiofacial syndrome (VCFS), a congenital disorder characterized by neural-crest-related developmental defects. In human and DGS models, *TBX1* haploinsufficiency causes craniofacial anomalies (Lindsay et al. 2001) and contributes to heart defects (Merscher et al. 2001). More precisely, the phenotype observed in the mutant mice for the T-box gene, *Tbx1*^+/-^, encompasses abnormal development of the skeletal structures derived from the first and second pharyngeal arches, with reduced dimension of the midface (Jerome and Papaioannou 2001); a similar situation found in the DS mouse models. In Dp(16)1Yey at an early stage (E11.5), *Ripply3* is overexpressed, and consecutively, *Tbx1* is downregulated in the midface precursor tissues. Still, we also detected a defect in cell proliferation of the NCC derivates in the first branchial arch, which also demonstrated a contribution to the midface shortening. In addition, our new model Dp(16)1Yey/*Ripply3^tm1b^* demonstrated an increased shape dimension in the structures corresponding to the midface compared to Dp(16)1Yey. Also, in some DS brain models, such as Ts1Rhr and Ts1Cje, the expression of the *Tbx1* gene was found to be significantly downregulated, and it might be involved in delayed fetal brain development and postnatal psychiatric phenotypes observed in DS (Shimizu et al. 2021). Considering this information, we postulated that the overexpression of *Ripply3* in DS mouse models will lead to a downregulation of *Tbx1*, in other DS organs and tissues, leading to additional changes. As such, some DS heart defects, such as the tetralogy of Fallot observed in some individuals, may be related to *Ripply3*-dependent downregulation of *Tbx1*, while in DGS, they are caused by the direct *Tbx1* haploinsufficiency (Merscher et al. 2001). Consequently, investigating treatment for DGS to reestablish a normal TBX1 function will also be of interest for Down syndrome, not only for the craniofacial but also for the brain and the heart function.

## Author contributions

Conceptualization: JTAS, ABZ, YH

Data Curation: JTAS, ABZ, YH

Formal Analysis: JTAS, CC, YH

Funding Acquisition: JTAS, ABZ, YH

Investigation: JTAS, CC, ABZ, YH

Methodology: JTAS, CC, YH

Project Administration: ABZ, YH

Supervision: JTAS, ABZ, YH

Validation: JTAS, ABZ, YH

Visualization: JTAS, ABZ, YH

Writing – Original Draft Preparation: JTAS, ABZ, YH

Writing – Review & Editing All

## Conflict of interest statement

The authors declare no competing interests.

## Funding

This work of the Interdisciplinary Thematic Institute IMCBio, as part of the ITI 2021-2028 program of the University of Strasbourg, CNRS and Inserm, was supported by IdEx Unistra (ANR-10-IDEX-0002), SFRI-STRAT’US project (ANR 20-SFRI-0012), INBS PHENOMIN (ANR-10-INBS-07) and EUR IMCBio (ANR-17-EURE-0023) under the framework of the French Investments for the Future Program. JT A received funding from the National Agency for Research and Development (ANID)/Scholarship Program/DOCTORADO BECAS CHILE/2020-72210028.

## Acknowledgments

We thank the Mouse Clinical Institute (PHENOMIN-ICS) for helping maintain the mutant mouse models. Special thanks to Loïc Lindner and Pauline Cayrou for helping with ddPCR design and training, to Sophie Brignon, Charley Pinault, and Aurélie Eisenmann at PHENOMIN-ICS and the IGBMC animal facility for their services, and to Patrick Reilly for proof-reading the manuscript.

## Data availability

The morphological data are deposited in Zenodo under the doi: 10.5281/zenodo.13639386

## Notes

https://doi.org/10.5281/zenodo.13639386

### Competing Interest Statement

The authors have declared no competing interest.

