## Supplementary information for "*Ripply3* overdosage induces mid-face shortening through *Tbx1* downregulation in Down syndrome models"

**SUPPLEMENTARY TABLES**

| **Landmarks Cranium** | |
|---|---|
| **1** | Nasale: Intersection of nasal bones, rostral point |
| **2** | Nasion: Intersection of nasal bones, caudal point |
| **3** | Bregma: intersection of frontal bones and parietal bones at midline |
| **4** | Intersection of parietal bones with anterior aspect of interparietal bone at midline |
| **5** | Intersection of interparietal bones with squamous portion of occipital bone at midline |
| **6** | Opisthion, midsagittal point on the posterior margin of the foramen magnum |
| **7** | Center of alveolar ridge over maxillary incisor, right side |
| **8** | Anterior Intersection of frontal process of maxilla with frontal bone, right side. |
| **9** | Anterior notch on frontal process lateral to infraorbital fissure, right side |
| **10** | Intersection of frontal process of maxilla with frontal and lacrimal bones, right side |
| **11** | Frontal-squasmosal intersection at temporal crest, right side |
| **12** | Intersection of zygoma (jugal) with zygomatic process of temporal, superior aspect, left side |
| **13** | Intersection of zygoma (jugal) with zygomatic process of temporal, inferior aspect, left side |
| **14** | Most posteroinferior point on the superior portion of the tympanic ring, right side |

| | |
|---|---|
| **15** | Center of alveolar ridge over maxillary incisor, left side |
| **16** | Anterior Intersection of frontal process of maxilla with frontal bone, left side. |
| **17** | Anterior notch on frontal process lateral to infraorbital fissure, left side |
| **18** | Intersection of frontal process of maxilla with frontal and lacrimal bones, left side |
| **19** | Frontal-squasmosal intersection at temporal crest, left side |
| **20** | Intersection of zygoma (jugal) with zygomatic process of temporal, superior aspect, right side |
| **21** | Intersection of zygoma (jugal) with zygomatic process of temporal, inferior aspect, right side |
| **22** | Most poteroinferior point on the superior portion of the tympanic ring, left side |
| **23** | Most anterior point of the anterior palatine foramen, right side |
| **24** | Most posterior point of the anterior palatine foramen, right side |
| **25** | Most infero lateral point on premaxilla-maxilla suture, right side |
| **26** | The anterior most point on the central ant/post axis of the right molar alveolus |
| **27** | Intersection of zygomatic process of maxilla with zygoma (jugal), inferior surface, right side |
| **28** | Lateral intersection of maxilla and palatine bone posterior to the third molar, right side |
| **29** | Joining of squasmosal body to zygomatic process of squasmosal, right side |
| **30** | Most inferior aspect of posterior tip of medial pterygoid process, right side |
| **31** | Most anterior point of the anterior palatine foramen, left side |
| **32** | Most posterior point of the anterior palatine foramen, left side |
| **33** | Most infero lateral point on premaxilla-maxilla suture, left side |
| **34** | The anterio most point on the central ant/post axis of the left molar alveolus |
| **35** | Intersection of zygomatic process of maxilla with zygoma (jugal), inferior surface, left side |
| **36** | Lateral intersection of maxilla and palatine bone posterior to the third molar, left side |
| **37** | Joining of squasmosal body to zygomatic process of squasmosal, left side |
| **38** | Most inferior aspect of posterior tip of medial pterygoid process, left side |
| **39** | Basion, midsagittal point on the anterior margin of the foramen magnum |

**Table S1:** 39 cranium Landmarks.

| | **Landmarks Mandible** |
|---|---|
| **1** | Apex of coronoid process, Right side |
| **2** | Intersection of molar alveolar rim and base of coronoid process, Right side |
| **3** | Anterior edge of alveolar process where first molar hits alveolus at the midline, Right side |
| **4** | Superior-most point on incisor alveolar rim at midline (at bone-tooth junctions), Right side |
| **5** | Inferior-most point on incisor alveolar rim at midline (at bone-tooth junction), Right side |
| **6** | Inferior point on mandibular symphysis, Right side |
| **7** | Anterior edge of the coalescence of curve of masseteric ridge with post-symphyseal rugged area, Right side |
| **8** | Tip of mandibular angle, Right side |

| | |
|---|---|
| **9** | Posterior midline point on condyle, Right side |
| **10** | Anterior midline point on condyle, Right side |
| **11** | Anterior edge of the mental foramen, Right side |
| **12** | Apex of the coronoid process, left side |
| **13** | Intersection of molar alveolar rim and base of coronoid process, left side |
| **14** | Anterior edge of alveolar process where first molar hits alveolus at the midline, left side |
| **15** | Superior-most point on incisor alveolar rim at midline (at bone-tooth junctions), left side |
| **16** | Inferior-most point on incisor alveolar rim at midline (at bone-tooth junction), left side |
| **17** | Inferior point on mandibular symphysis, left side |
| **18** | Anterior edge of the coalescence of curve of masseteric ridge with post-symphyseal rugged area, left side |
| **19** | Tip of mandibular angle, left side |
| **20** | Posterior midline point on condyle, left side |
| **21** | Anterior midline point on condyle, left side |
| **22** | Anterior edge of the mental foramen, left side |

**Table S2:** 22 mandible Landmarks.

### SUPPLEMENTARY FIGURES

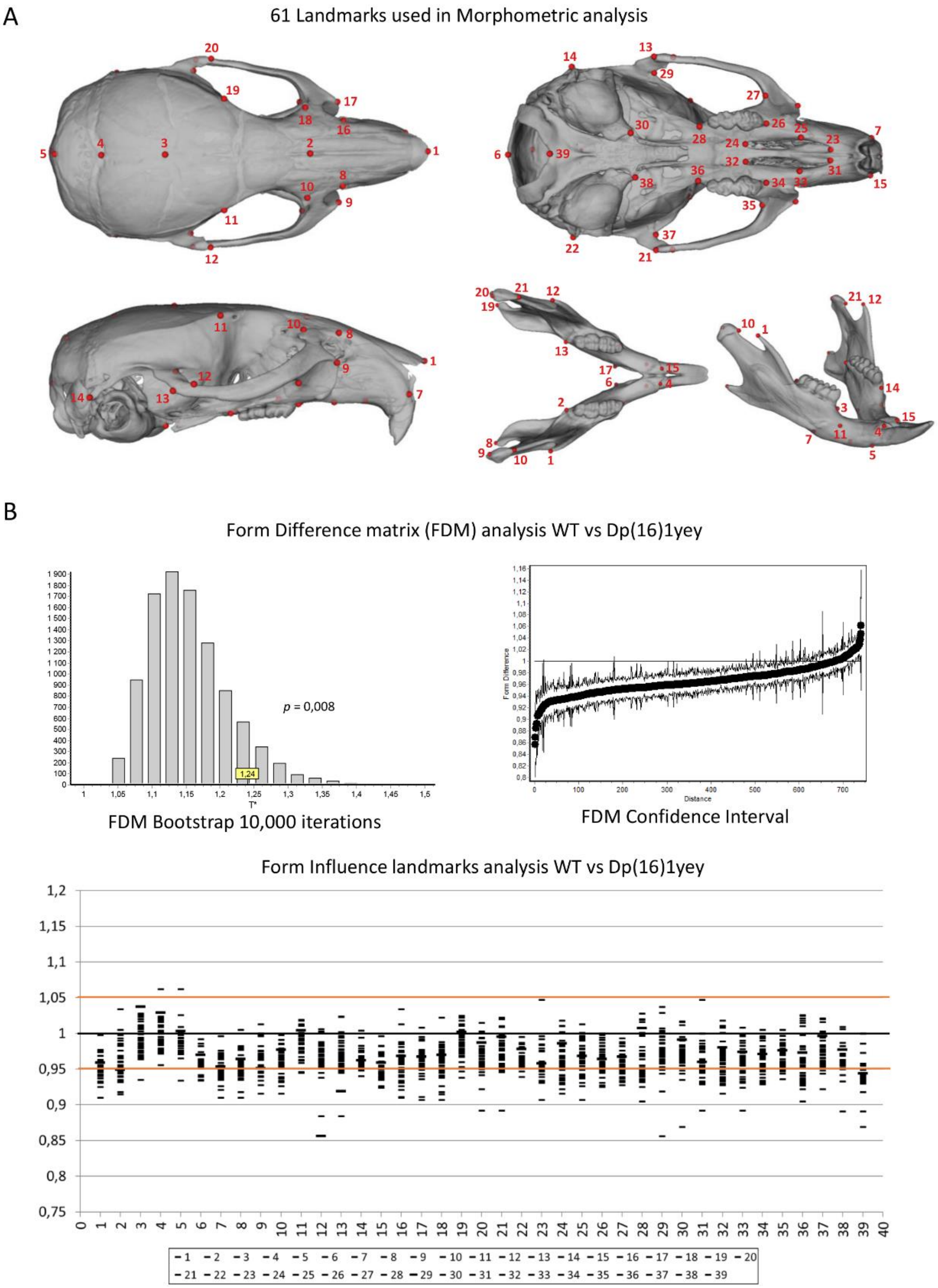


**Figure S1. Landmark detailed analysis of the cranium and mandibular phenotypes in Dp(16)1Yey. (A)** 61 Landmarks used in the CF analysis of Dp(16)1Yey (and in all the other DS models). 39 in the cranium and 22 in the mandible. **(B)** Form difference matrix analysis: FDM Bootstrap with 10,000 iterations showing significant changes in Form (p=0.008). The FDM confidence interval graph shows a decrease of more than 90% of the distances measured. Form Influence landmarks graphic, showing the landmarks that present a relative Euclidean distance > 1.05 or < 0.95 (outside of the confidence interval 97,8%, red lines) and a general reduction of all dimensions.

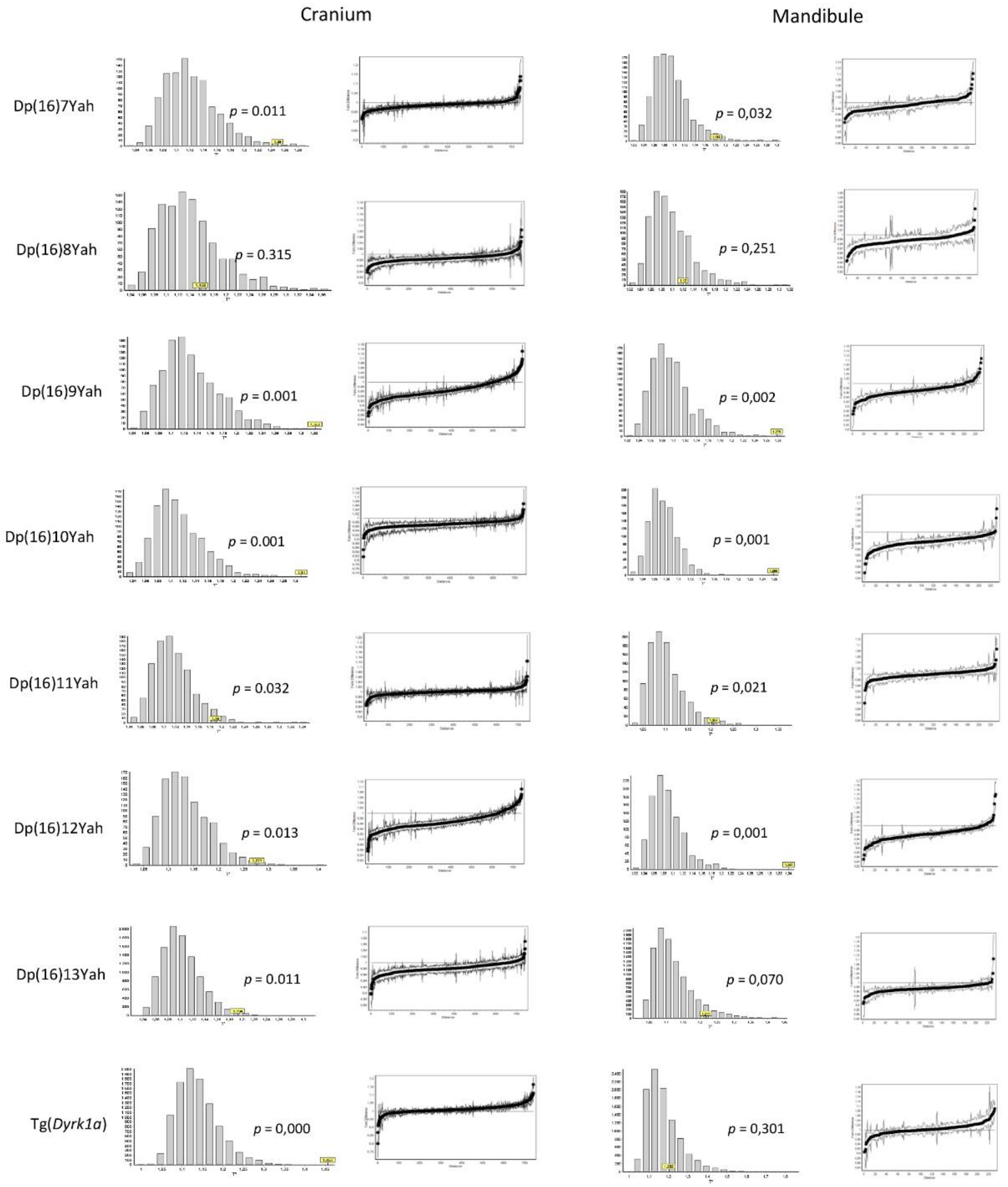


**Figure S2. Form difference matrix analysis of the new panel of mouse models, plus Tg(Dyrk1a).** On the left, Cranium analysis with from difference matrix (FDM) tested for with Bootstrap (graph) showing significant changes in form for all the models with form difference interval graph, except Dp(16)8Yah (p=0.315). On the right, the same graphs for the mandible show significant changes in form for all the models except Dp(16)8Yah (p=0.215). On the right, the FDM confidence interval graph shows the ratio of the distances measured.

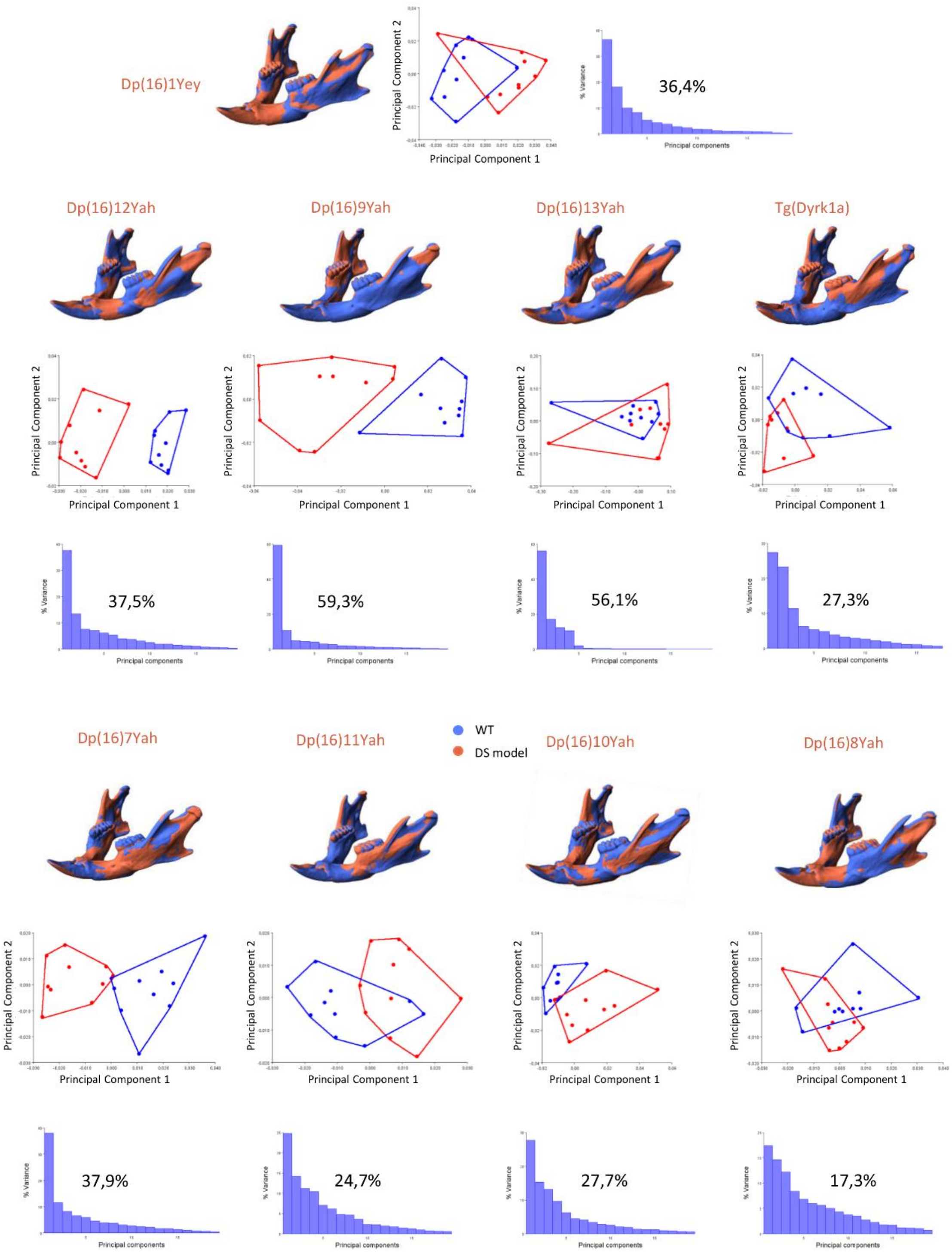


**Figure S3. New DS mouse models mapping the location of dosage-sensitive genes that cause the craniofacial dysmorphology of Dp(16)1Yey (Mandibles).** Morphometric analysis of the mandibles of the new panel of DS mouse models, plus Tg(Dyrk1a) and Dp(16)1yey. Shape difference warping to display the mandible parts with decreased dimensions in DS models (in blue), and with increased dimensions (in red). PCA (first two components) of general Procrustes analysis of aligned mandible shapes for every model with the contribution to the explained variance for each dimension.

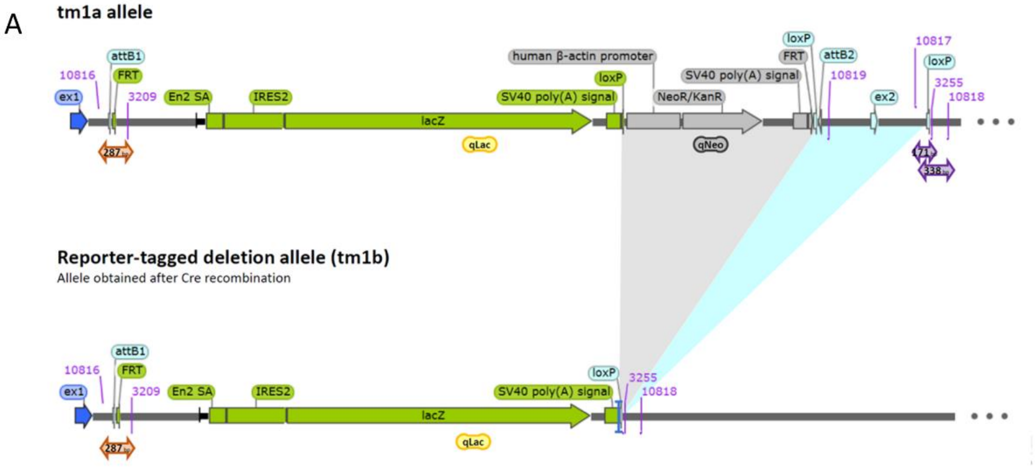


**Figure S4. Schematic representation of the *Ripply3$^{tm1b}$* knock-out model** derived from *Ripply3$^{tm1a}$*.

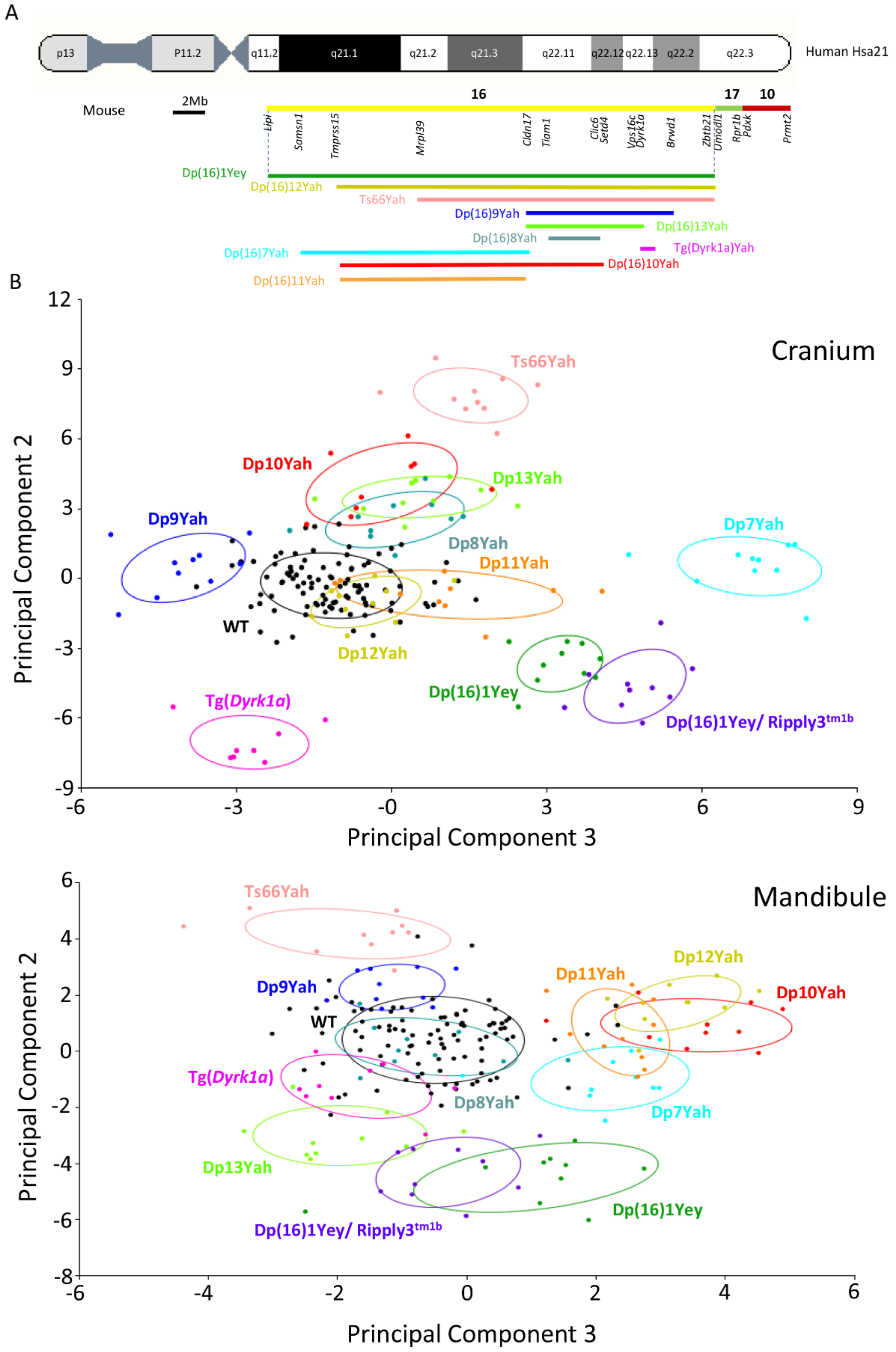


**Figure S5. PC2 and PC3 projection of the CF analysis discriminate the genotypes of DS models. (A)** Schematic representation of DS models and their relative position to HSA21. **(B)** PC2 and PC3 graphs for the cranium and mandible. Integrative multivariate analysis of all the models used in this study, plus Ts66Yah, vs wild-type. The PCAs (PC2 vs PC3) correspond to a canonical variate analysis (Procrustes Distance Multiple Permutations test at 1000 iterations). DS strains show significant differences compared with their wild-type controls. DS models were separated into four main groups in the cranium graph, as in PC1, but with different distributions. For the mandible, the graph showed a primary group of 5 models close to the wild-type and two branches separated on PC2, as observed in PC1 vs PC2.
